# A genomic toolkit for surveillance and elimination of the principal malaria vectors in Southeast Asia

**DOI:** 10.64898/2025.12.04.691712

**Authors:** Tristan P. W. Dennis, Amelie Vantaux, Kevin C. Kobylinski, Mohammad Shaiful Alam, Benoit Witkowski, Hasan Mohammad Al-Amin, Neil F. Lobo, Ayley J. Wilson, Eleanor Drury, Sonia Goncalves, Nick Harding, Christopher S. Clarkson, Anastasia Hernandez-Koutoucheva, Dominic Kwiatkowski, Alistair Miles, Martin J. Donnelly, Brandyce St. Laurent

## Abstract

While substantial progress has been made toward malaria elimination in Southeast Asia, major challenges remain. The principal mosquito vectors, *Anopheles dirus* and *Anopheles minimus*, inhabit diverse ecological niches and exhibit a wide range of behavioural and physiological insecticide resistance phenotypes, complicating vector control efforts. In Africa, genomic surveillance has transformed our understanding of vector evolution and resistance, supported by open-access tools and data. Comparable resources for Southeast Asian vectors remain limited.

Here, we release Adir1.0, a curated, analysis-ready catalog of 540 *An. dirus* whole-genome sequences, accessible for interactive cloud-based analysis via the malariagen-data-python application programming interface (API). Alongside, we provide an updated Amin1.0 API to enhance functionality and usability, enabling integrated analyses across historical and new datasets.

We demonstrate these resources by performing the first exploratory population genomic analysis of *An. dirus* from Bangladesh, Thailand, and Cambodia, revealing population structure, candidate regions of physiological resistance, and genomic structural variation. Together, these resources provide a foundation for genomic surveillance studies to inform vector control and malaria elimination strategies in Southeast Asia.

## Introduction

Southeast Asia accounts for approximately 5 million malaria cases annually, but is making substantial progress toward elimination. Major challenges threaten this progress including parasite drug resistance^1–3^, emerging behavioural and physiological insecticide resistance in vectors^4,5,6^, and rapidly changing land-use patterns that alter vector ecology^7,8^ and bring human and vector populations into closer contact. These pressures are compounded by risks of parasite reintroduction and zoonotic transmission^9^.

Deforestation and urbanization are fragmenting Southeast Asian landscapes, creating heterogeneous habitats that favour different vector species. The two dominant vector species complexes, *Anopheles dirus* and *An. minimus*, occupy distinct ecological niches with divergent insecticide resistance profiles. *An. minimus* thrives in human-modified environments such as rice paddies, where insecticide exposure may have driven shifts toward outdoor biting behavior (exophily) and increasing insecticide resistance in some populations^10^. In contrast, *An. dirus*, the region’s most important vector, is strongly associated with forest habitats, bites outdoors and in the early evening^11^, and drives malaria transmission in forest workers and other malaria-naïve populations. The *An. dirus* complex comprises multiple taxa, including *An. dirus* sensu stricto (*An. dirus* s.s.) predominant in Vietnam, Laos and Cambodia, and *An. baimaii* endemic to Bangladesh, Myanmar and western Thailand^12,13^. Whilst these are the primary vector species there is an unusually high diversity in vectors in the region many of which are understudied compared to African mosquito species. In Thailand, for example, there are approximately 20 Anopheles species which are human malaria vectors^14^.

Genomic approaches are essential for resolving the true complexity of Southeast Asian vector biology and informing vector control. Preliminary morphological and molecular work has delineated subspecies within the *An. dirus* complex^12,13^, but the extent of reproductive isolation, gene flow, and cryptic speciation remains unclear. Experience with other malaria vectors demonstrates that only genome-scale data can reveal this hidden complexity: *An. gambiae* ss and *An. coluzzii* are morphologically indistinguishable, and only genome-wide sequencing^15^ was able to determine that these are distinct species with different epidemiological roles^16^. Similarly, our previous work in the genomic surveillance of *An. minimus* uncovered intricate population structure, potential cryptic species, and selection signatures at insecticide resistance loci^6^. For *An. dirus*, these insights are operationally critical. As landscapes fragment and mosquito populations encounter intensifying insecticide exposure from agriculture and vector control, genomic surveillance enables early detection of resistance alleles^17^ and mapping of gene flow that shapes resistance spread, as well as identifying important hotspots for transmission. Furthermore, as Southeast Asia approaches malaria elimination and the cost incurred per case averted grows dramatically, genomic surveillance data can help guide novel intervention deployment, including treated hammocks, repellents, and baited traps, in forest-fringe and other hard to reach communities. Coordinated genomic surveillance of malaria parasites in the Greater Mekong Subregion provides a template of how molecular data can generate actionable intelligence for elimination strategies^1,18^.

Vector genomic surveillance is a key tool in the arsenal against malaria and is transforming the evidence base for malaria vector control. The Malaria Vector Genome Observatory (VObs), is a collaborative project developing capacity for *Anopheles* genomics, and has led the development of cloud-native data and analysis tools to promote open data sharing, co-operation and capacity building^19^. By combining capacity building with publicly accessible genomic data, VObs has facilitated landmark synthesis studies of *An. gambiae*^*20,21*^, *An. funestus*^*22*^, and characterised insecticide resistance mechanisms and vector invasions^23^. Critically, preliminary genomic surveillance of *An. minimus* in Southeast Asia by VObs^6^ has revealed complex population structure and insecticide resistance-associated loci.

Here we present a genomic toolkit to facilitate genomic surveillance of *An. dirus* in Southeast Asia. We sequence the genomes of 540 *An. dirus s*.*l*. from Bangladesh, Thailand and Cambodia, and present the results of population genetic analysis characterising their population structure and diversity, and performing a survey for potential insecticide resistance determinants. These data are made publicly available in association with the malariagen_data Python Application Programming Interface (API)^19^, which facilitates easy public access and interactive analysis. We also provide an updated version of the previously released Amin1 *An. minimus* API and data resource. We hope these data and tools will form the foundation of further genomic surveillance studies of malaria vectors in Southeast Asia.

## Results

### Data and sequencing

We sequenced the genomes of 540 *An. dirus* complex samples from Bangladesh (n=47), Cambodia (n=247) and Thailand (n=219) (**Figure 1A**) to a target coverage of 30X (see *Methods*). The mean depth-of-coverage varies between 8 and 121, with a median of 35.

**Figure 1.**
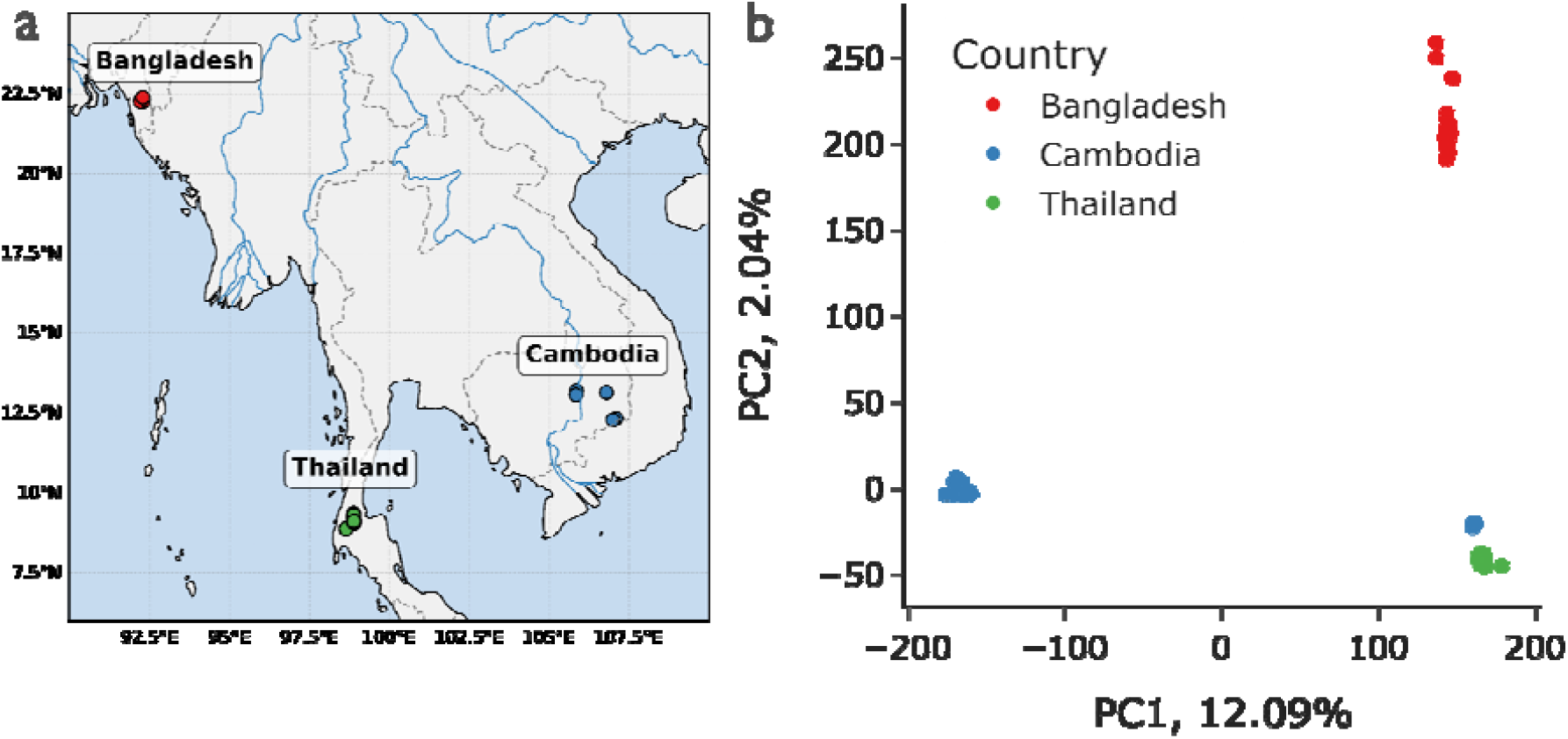
Sampling locations and population structure of *An. dirus* s.l. **a)** Sampling locations. X and Y axis denote latitude and longitude respectively. Sampling locations are marked by points. **b)** Principal component analysis (PCA) of *An. dirus s*.*l*., based on 50,000 randomly chosen variants from the longest genomic contig (KB672490). Samples are coloured by country of collection. Cambodian samples on the left of PC1 are *An. dirus*, the remainder are *An. baimaii* from Bangladesh, Thailand and a single location in Cambodia.

### Open data and application programming interface (API)

These data were sequenced and analysed as a member project of the Vector Genome Observatory - Asia (VObs-Asia). The population genomic data generated in these projects will help us understand the dynamics of malaria transmission in Asia, and improve malaria elimination. As part of VObs, SNP data are routinely released in publicly accessible cloud storage buckets. These data can be easily accessed and analysed using the open-source malariagen_data Python package. malariagen_data provides a suite of functions (application programming interface, or API) for analysing mosquito vector genome data, and can be used in the cloud, using free Google Colaboratory instances, or on a local machine. Here, we present the *Anopheles dirus Phase 1*.*0* (*Adir1*.*0)* data release (https://malariagen.github.io/vector-data/adir1/adir1.0.html) and accompanying API (https://malariagen.github.io/malariagen-data-python/latest/Adir1.html), as well as an updated version of the previously released *Amin1* data (https://malariagen.github.io/vector-data/amin1/intro.html) and API release (https://malariagen.github.io/malariagen-data-python/latest/Amin1.html). We use the *Adir1* data release, and malariagen_data API to produce a set of initial population genomic analyses of *An. dirus* in Bangladesh, Thailand and Cambodia, as well as additional analysis of *An. minimus*, building on our previous work^6^. These analyses serve as a demonstration of the tools and data, as well as the first large-scale set of population genomic analyses of *An. dirus*.

### Genetic structure and diversity of An. dirus

The *An. dirus* species complex contains multiple closely related members. Two of these, *An. baimaii and An. dirus sensu stricto* (previously called An. dirus D), and *An. dirus A* respectively, are present in our dataset (**Table S1**). From now on, *An. dirus s*.*l*. will refer to the *An. dirus* complex (*sensu lato*), *An. dirus* will refer to *An. dirus sensu stricto*, and *An. baimaii* will refer to *An. dirus baimaii*. Principal component analysis (PCA) of genotypes from the longest contig in the genome (KB672490) reveals three major groups. (**Figure 1B**). PC1, which explains 12.09% of the variation in the data, separates *An. baimaii* and *An. dirus*, with a subgroup of samples initially identified as *An. dirus* from a single site in Cambodia grouping with *An. baimaii* samples from Cambodia (**Figure 1B**). The third main cluster contains *An. dirus ss* samples from Cambodia (**Figure 1B**). PCA groupings are used to define analysis cohorts subsequently.

### Genetic diversity

We estimated genetic diversity in *An. dirus* s.l. by estimating the count, and fraction of an individual sample’s genome in runs of homozygosity (n/fROH). fROH, which approximates the individual inbreeding coefficient^26^, is similar within and between populations (mostly below 5% (**Figure 2A**)), with Cambodian/Thai *An. baimaii*, tending towards a greater fROH per-sample. nROH varied more, with Cambodian/Thai *An. baimaii* having a greater nROH than Cambodian *An. dirus*. In general, these signatures are similar to the highly diverse, connected populations of *An. gambiae*^*20*^ and *An. funestus*^*27*^, and endemic (Pakistan/Afghanistan) populations of *An. stephensi* in Asia^23^, as opposed to more isolated *An. funestus* populations, or populations of *An. gambiae* and invasive *An. stephensi* (Sudan) that have undergone recent bottlenecks as a result of vector control or recent invasions^20,23^. We examined nucleotide diversity (π) in *An. dirus, An. minimus*, compared to comparator populations of *An. gambiae, An. funestus* and *An. stephensi*. π was highest in Bangladesh *An. baimaii -* consistent with the ROH data (**Figure 2B**). π from the other cohorts was similar to *An. minimus* from Cambodia^6^ and was higher than both endemic and invasive *An. stephensi*. π in *An. funestus* and *An. gambiae* from Tanzania was higher than all cohorts included in this analysis (**FIgure 2B**). Tajima’s D was consistently negative across all *An. dirus* cohorts (**Figure 2B**), with Cambodian/Thai *An. baimaii* showing the most negative values. These values were similar across *An. dirus, An. minimus*, endemic *An. stephensi, An. funestus* and *An. gambiae*.

**Figure 2:**
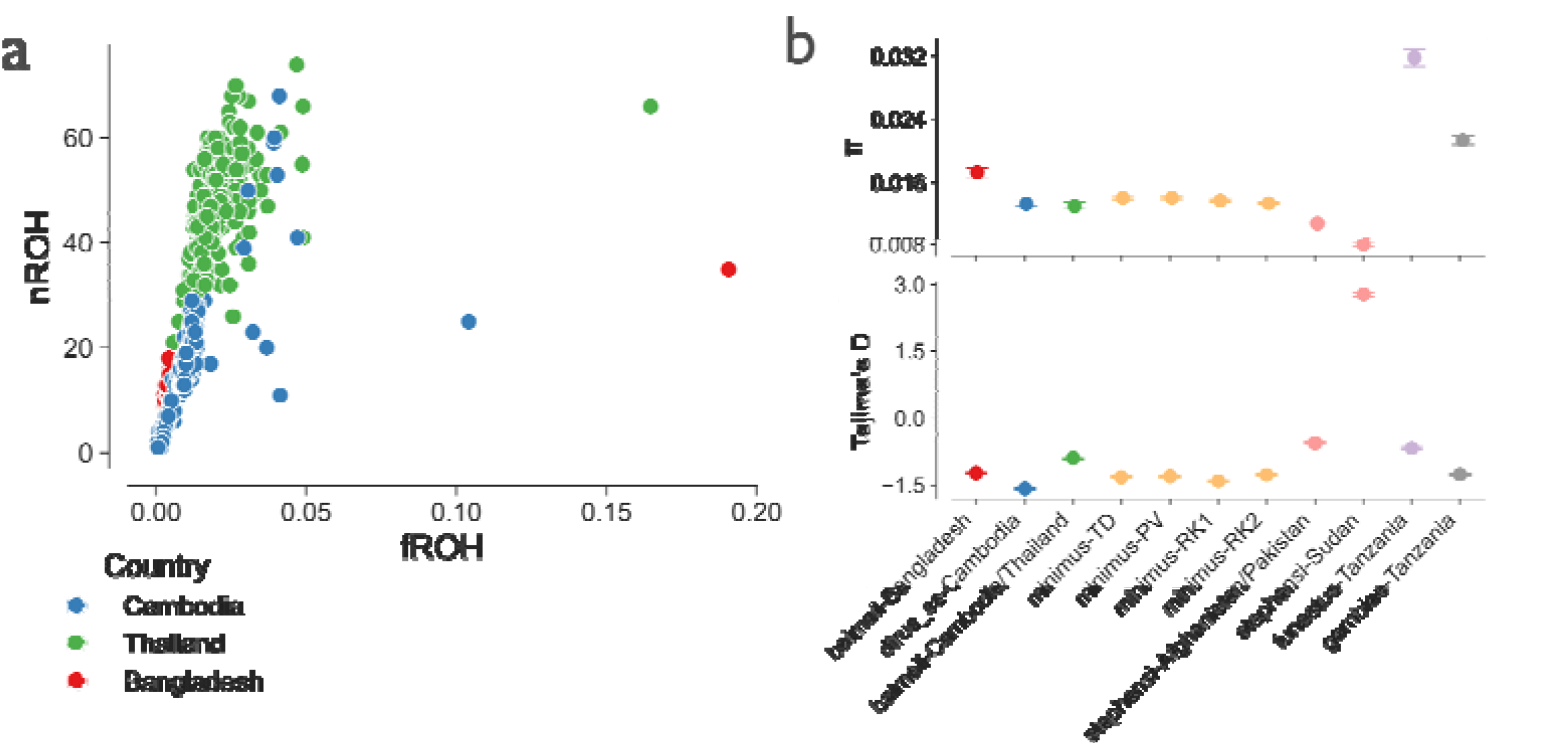
Genetic diversity of *An. dirus s*.*l*. (a)shows the joint distribution of the total fraction of an individual genome in runs of homozygosity (ROH) (x axis), and the number of ROH per-individual (y axis). Points indicate individual samples and are coloured by country of collection as per the PCA in **Figure 1B. (b)** and **(c)** show median nucleotide diversity (π) and Tajima’s D respectively, calculated in windows genomewide for three populations delineated in the PCA above (e.g. Cambodian *An. baimaii* is grouped with Thailand *baimaii*). Also included are comparator populations of other species in the Vector Observatory: Cambodian *An. minimus* (broken down by population^6^), Sudanese and Afghanistan/Pakistan *An. stephensi*^*23*^, and Tanzanian *An. funestus*^*27*^ / *An. gambiae*^*20*^. Error bars indicate 95% confidence intervals estimated by block-jackknife. Point colour for all plots denotes groups of samples broken down by taxon and country as per the PCA in **Figure 1B** (see legend in **(a)**).

### Genome-wide differentiation and chromosomal inversions

Genomewide differentiation (*F*_*st*_) was high between *An. baimaii* and *An. dirus* (**Figure 3A**) (median 0.55 and 0.52 between Cambodian/Thai *An. baimaii*, Bangladeshi *An. baimaii*, and Cambodian *An. dirus*, respectively), and far lower between the *An. baimaii* populations (median 0.05). Areas of elevated differentiation were pronounced across the whole or parts of some contigs, with a tendency toward elevated *F*_*st*_ in shorter contigs. Previous work has identified five inversion polymorphisms occurring across chromosomes 2, 3 and X^28–31^, and a linkage map has been produced, scaffolding the inversions across the two autosomes and the sex chromosome. We investigated variation in population structure across the genome using windowed PCA - an approach used to detect inversion polymorphisms in other *Anopheles* taxa^22,23^. (**Figure 4**). We found evidence of several inversion polymorphisms. While *An. dirus* appeared homosequential (**Figure S1**), *An. baimaii* had several signatures of at least four large inversion polymorphisms, including three on the autosomes^30^, and one on the X chromosome. Each population showed different karyotypes. Cambodian *An. baimaii* was largely homosequential, with the exception of a small region (contig KB673534, KB672846 and KB672991 in two karyotypes, also shared with Bangladesh and Thailand (which had three karyotypes). The largest inversion spanned contigs KB673090 and KB673201, was present in Bangladesh and Thailand, which possessed distinct karyotypes. Finally, Bangladesh and Thailand shared karyotypes of a smaller inversion present on contigs KB672868, KB672713 and KB673645 (**Figure 4**). Bangladesh also had a possible inversion present on KB672491 and KB672824, as well as a large region of contigs KB672490, KB672902, KB672791, KB672602 and KB672835 that showed various outgroups, but not the banding pattern characteristic of inversion polymorphisms (**Figure 4**).

**Figure 3.**
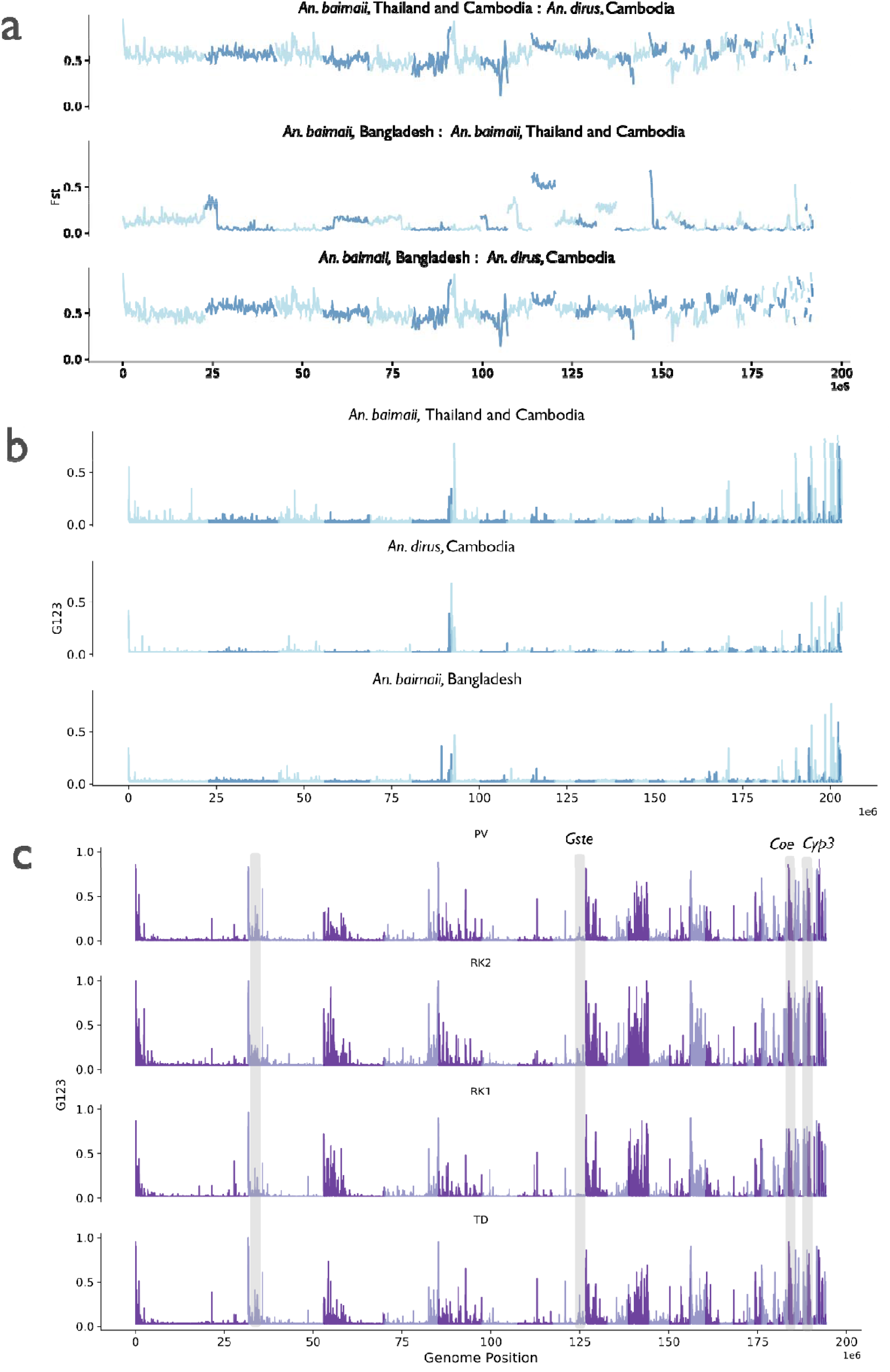
(previous page): Genome-wide selection scans. **a**) shows *F*_*st*_, and **b**) shows the G123 statistic^56^ for *An. dirus*. Statistics were calculated in windows per-contig, and plotted by contig in size order, descending. Alternating shades on the lines indicate alternating contigs. The x-axis, shared between all plots, indicates cumulative genomic position calculated across all contigs. The y-axes denote *F*_*st*_ (a) and G123 (b) respectively. Plots are panelled by between-population comparison (a) and population (b), respectively, and are labelled accordingly above each subplot. **b)** shows the G123 statistic calculated for *An. minimus*, with sweeps at resistance-associated loci shaded in grey and labelled above. Note genome assemblies differ between *An. dirus* and *An. minimus*, so genomic positions on the x axis differ between **a/b** and **c**.

**Figure 4.**
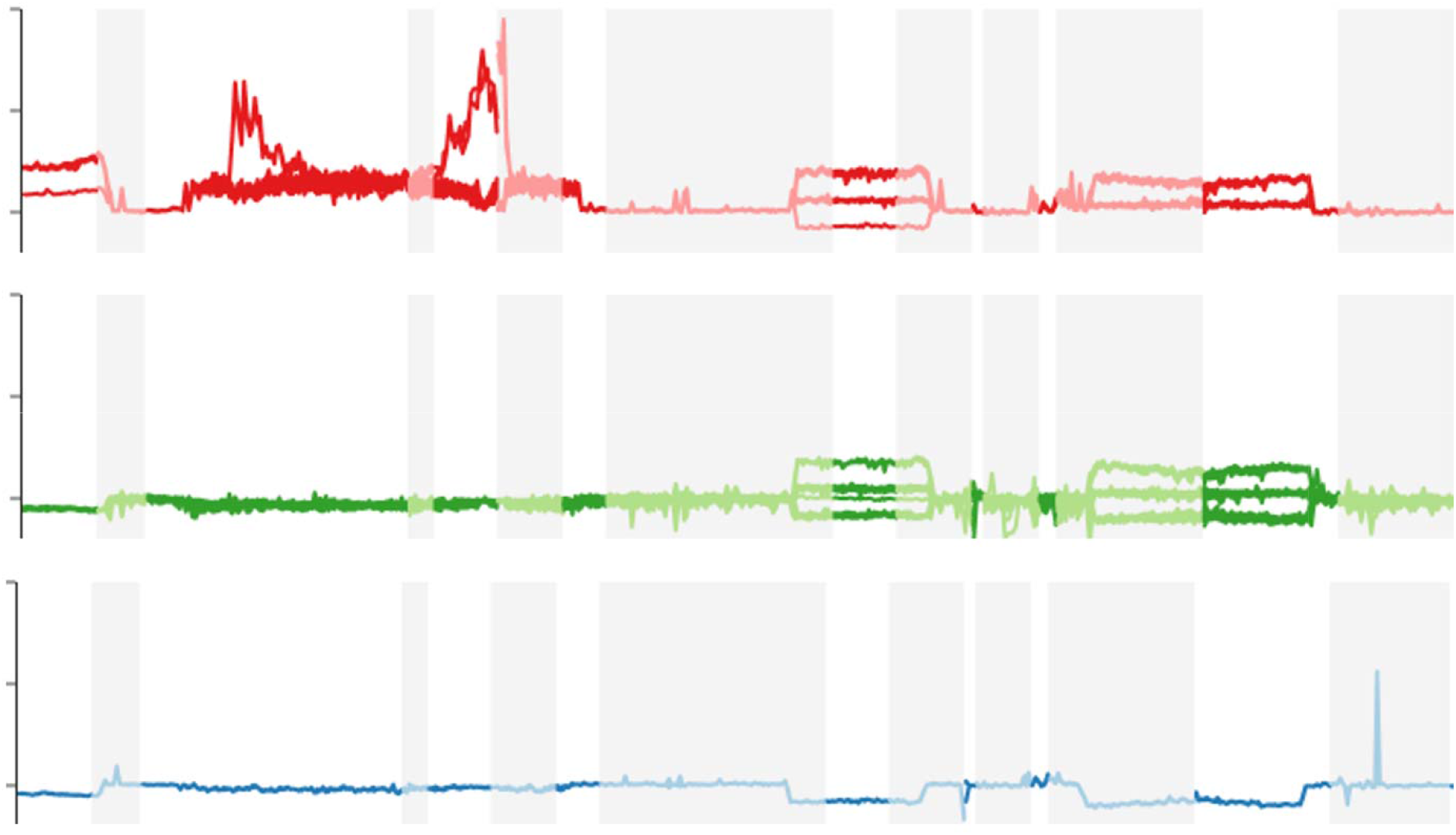
Windowed PCA by contig for *An. baimaii*. PCA was calculated in 100,000 BP windows across each contig, and PC1 (y axis) was plotted for each sample. Rows and colours denote country of collection (red: Bangladesh, green: Thailand, blue: Cambodia) along genomic position (x axis), for each contig in the *An. dirus* reference genome assembly AdirW1. Different contigs are shown by white/light grey shading in the plot background, and labelled below the x axis.

### Selective sweeps

Estimation of selection within species/populations using the G123 statistic (**Figure 3B)** showed that a number of regions of the genome exhibited signatures characteristic of selection. As in the interspecific pairwise *F*_*st*_ analysis (**Figure 3A**) there was a tendency for elevated G123 values to occur in shorter contigs. Given in other *Anopheles* species selection signatures are commonly reflective of recent adaptation to insecticide pressure^17,32^ we preferentially examined the areas under the outlier peaks for genes or gene families associated with insecticide resistance in other *Anopheles* vector taxa (e.g. at the *Voltage gated sodium channel (Vgsc)* or the *Cytochrome p450 subfamily 6 (Cyp6)* cluster)^23,33,34^ **(Table S2)**. No loci commonly implicated in the evolution of insecticide resistance were found within the swept regions. However we did identify a number of loci with swept regions from detoxification gene families which could conceivably be driving the selective sweeps, although additional validation is clearly required. We found very elevated G123 (0.7) in a small contig (KB673079) in all three populations, in a window centred on the *Glutathione-S-transferase subfamily theta 1 (Gstt1)* gene. In other anophelines, glutathione-S-transferases (*Gste*) are major determinants of metabolic resistance^35^. Additionally, we found evidence of elevated G123 (0.7) at a carboxylesterase (*Coe120*) in Cambodia/Thai *An. baimaii*, and Bangladesh *An. baimaii*. A smaller sweep (G123 of 0.1) at the *Cyp6AJ1* was also present in Cambodia/Thai *An. baimaii*. The largest peaks appeared at the ends of contigs, or in small contigs, suggesting regions that were hard to assemble (e.g. due to low accessibility, presence of structural variation, repetitive regions or centromeres) may have caused this signal by artificially inflating homozygosity. For comparison, we also calculated G123 for the four Cambodian *An. minimus* populations we analysed previously^6^. In contrast to *An. dirus*, we found a large number of genomic regions in elevated G123. However, most of these were also at the end of contigs, or contained un-annotated genes. By preferentially searching outlier peaks for gene families implicated in resistance in other *Anopheles* taxa, we found evidence of elevated G123 at the *Gste* and *UDP-glucose:glycoprotein glucosyltransferase (UGT)* loci^36^ (consistent with elevated *Fst* signals in our previous work^6^). We also found evidence of sweeps at a cluster of *Cyp450* genes orthologous to the *Cyp3* family in *An. gambiae*, and a pair of *Carboxylesterase* genes (**Table S3**). These peaks were present in all four cohorts.

### Insecticide resistance mutations

We searched in a set of genes known to be involved in target-site resistance in *An. gambiae* for non-synonymous mutations implicated in resistance. We searched the *Voltage-gated sodium channel* (*Vgsc*), *Acetylcholinesterase (Ace1), Resistance to dieldrin (*Rdl) genes. We also searched the region containing the *Glutamate receptor, anionic (GluCl)* - implicated in resistance to Ivermectin, under trial for vector control in the region^37^. We found that these genes were surprisingly monomorphic, with no mutations segregating at >10% in any cohort in *GluCl*, and *Rdl*, 5 in the *Vgsc* and 8 in *Ace1* (Table S3). Sequence comparison in multiple sequence alignments confirmed that none of the non-synonymous mutations identified corresponded to the L995F, V402L/I527M mutations associated with resistance in *Vgsc* in *An. gambiae*, or the *G280S* mutation conferring resistance in *Ace1* in *An. gambiae*. (**Table S4**).

## Discussion

Here, we present a genomic toolkit that will form the foundation of effective genomic surveillance of malaria mosquitoes in Southeast Asia: an initial dataset of 540 *An. dirus s*.*l*. mosquito genomes, accompanied with the malariagen_data Python API, facilitating free access and analysis of these data. Moreover, we provide a substantial update to the functionality of the *Amin1* API, facilitating further analysis of this malaria vector. Our work provides a number of interesting findings that will form the basis of future work interrogating the biology and evolution of major malaria vectors in Southeast Asia.

Physiological resistance is reportedly scarce in *An. dirus*, and sporadically evolves in *An. minimus*^*5,38,39*^. Our findings in *An. dirus* further support this, with only weak evidence for selection at two gene families (*Glutathione-s-transferase theta*, and *Carboxylesterase*) sharing a close functional relationship with the *Glutathione-s-transferase epsilon* and *Carboxylesterase alpha-esterase* determinants of resistance in other mosquito species, as well as a weak sweep at a *Cyp6* locus. Aside, we found no evidence for selection, or appreciable frequencies of possible target-site resistance mutations, in gene families that underpin resistance in other Asian and African^5,23,33,40–44^ vectors. This is in contrast to *An. minimus*, where we found a substantial number of genome-wide selective sweeps - mostly containing un-annotated genes, including a number containing possible resistance determinants (e.g. *Cyp3*), confirming our previous work where we found elevated *F*_*st*_ at the *Gste2* and *U-ggst* detoxification loci. These differences could reflect the different life-histories of each species: *An. dirus* live in deep forest habitats and forest-adjacent plantations (e.g. rubber plantations) and are are exophilic. *An. dirus* will have had less contact with home-based control measures such as bednets and indoor residual sprays, and will be less prone to contamination of larval sites with agricultural pesticides. By contrast, *An. minimus* lives in peri-domestic and agricultural habitats, and has shown both endo- and exophilic behaviour^10,11,45,46^.

An. dirus populations maintain relatively high genetic diversity, with ROH signatures consistent with large, well-connected populations. The elevated fROH and nROH in Cambodian/Thai An. baimaii, coupled with strongly negative Tajima’s D values, may reflect range-edge effects or recent expansion, contrasting with the higher diversity observed in Bangladesh *An. baimaii*. Our windowed PCA analysis confirms previous observations of high levels of inversion polymorphism in *An. baimaii*^*28,30,31*^, while *An. dirus* remained largely homosequential. Within *An. baimaii* Cambodian samples were predominantly homosequential, but Bangladesh and Thailand harbored multiple inversions with population-specific karyotypes. This chromosomal heterogeneity may reflect local adaptation to the diversity of ecological conditions in the region. For genomic surveillance, these inversions are critical as they suppress recombination and create linkage blocks that may harbor clusters of adaptive alleles, including potential resistance determinants^47^.

Enhanced surveillance of mosquito vector populations is essential for understanding the complexity of malaria transmission scenarios in Southeast Asia^48^. The incredible ability of mosquito vectors to adapt to interventions and changing landscapes raises a number of challenges, such as physiological insecticide resistance, behavioural adaptations and changes in species composition. All of these factors have implications for planning optimal malaria control as the region transitions to malaria elimination. Genomic surveillance has a key role to play during this period of change: understanding the extent to which mosquito vectors represent different species, surveilling for evolutionary threats like insecticide resistance, and providing more general intuition as to the major mechanisms of adaptation to control in mosquito populations. Future work, including expanding the scale of sampling and the addition of phased haplotypes and copy-number variant calls, will facilitate more sophisticated analyses into the mechanisms of adaptation and resistance, species delineation, and landscape genetics for identifying population sources and sinks. Together, the Adir1 data and API release, combined with the updated Amin1 resource, position the malaria vector community to conduct coordinated genomic surveillance across Southeast Asia, paralleling the successful genomic surveillance of parasites in the Greater Mekong Subregion^1^ and providing actionable intelligence to guide elimination strategies in this rapidly changing region.

## Methods

### Sample collection and processing

Adult *An. baimaii* were collected in Bangladesh (n=47) in 2018 using a combination of light traps and human-landing catches, and in Thailand (n=219) between February and October 2019 using human-landing catches^49^. *An. dirus sensu stricto* were collected in Cambodia between October 2017 and November 2020 (n=274) using human-baited double tent traps. DNA was extracted using a QIAGEN DNeasy blood and tissue kit. For more per-sample information (e.g. specific location or date), see **Table S1**.

### Whole genome sequencing and analysis

The samples were processed as part of the MalariaGEN Vector Observatory Asia (VObs-Asia) *Anopheles* genomic surveillance project [https://www.malariagen.net/project/vector-observatory-asia/]. Briefly, the mosquitoes were individually whole-genome-sequenced on an Illumina NovaSeq 6000s instrument. Reads were aligned to the *An. dirus* reference genome AdirW1^50^ with Burrows-Wheeler Aligner (BWA) version v0.7.15^51^. Indel realignment was performed using Genome Analysis Toolkit (GATK) version 3.7-0 RealignerTargetCreator and IndelRealigner^52^. Single nucleotide polymorphisms were called using GATK version 3.7-0 UnifiedGenotyper. Genotypes were called for each sample independently, in genotyping mode, given all possible alleles at all genomic sites where the reference base was not “N”.

Complete specifications of the alignment and genotyping pipelines are available from the malariagen/pipelines GitHub repository [https://github.com/malariagen/pipelines/]. The aligned sequences in BAM format were stored in the European Nucleotide Archive (ENA).

The identification of high-quality SNPs and haplotypes were conducted using BWA version 0.7.15 and GATK version 3.7-0. Quality control involved removal of samples with low mean coverage, removing cross-contaminated samples and running PCA to identify and remove population outliers. Full quality control methods are available on the MalariaGEN vector data user guide [https://malariagen.github.io/vector-data/ag3/methods.html].

We used a hard filter to identify genomic sites where SNP calling and genotyping is likely to be less reliable. More information on site filters can be found on the MalariaGEN vector data partner user guide for Adir1.0^24^.

### Population genetics

All population genetics was performed using the malariagen_data Python API. All code and detailed parameters can be found in Jupyter notebooks at https://github.com/tristanpwdennis/sea_vector/. Plotting was performed using plotly^53^, seaborn^54^, and cartopy^55^. Analyses are fully reproducible using the malariagen_data API.

Principal component analysis was performed using 100,000 randomly chosen SNPs from the largest contig (KB672490). Genomewide selection scans (GWSS) were performed using windowed *F*_*st*_, and G123^56^, with window sizes of 100 and 2kb respectively, on analysis groupings defined in the PCA. Runs of homozygosity (ROH) were inferred per-sample, with a length cutoff of 100,000 bp applied, to distinguish between regions of the genome homozygous as a result of linkage disequilibrium vs recent inbreeding^26^. Thetas (π and Tajima’s D) were calculated using 200 block-jacknife resampling replicates, whereby for the whole genome, a block was deleted, and thetas recalculated, providing estimates of the 95% confidence interval. Genome-wide *F*_*st*_ was calculated in sliding windows of 50,000 bp. G123 was calculated in windows of 2000bp. For all GWSS, samples were grouped according to the PCA results (**Figure 1b, *Results***). Outlier peaks were identified by taking the 95% quantile of G123 for each population and cutting off windows that fell below that threshold. Sliding-window PCA was calculated per-contig in windows of 50,000bp.

## Data availability

Code supporting this study can be found at https://github.com/tristanpwdennis/sea_vector/. Catalogs of raw read, alignment, and variant call data, and instructions for download, can be found at: https://malariagen.github.io/vector-data/adir1/adir1.0.html and https://malariagen.github.io/vector-data/amin1/intro.html.

## Acknowledgements

We thank the following personnel in Thailand for supporting field collections of *An. dirus* in Surat Thani: Narenrit Wamaket, Jeeraphat Sirichaisinthop, Veerast Suwan and the Entomology Team at the Surat Thani Vector-Borne Diseases Control Center. We also thank Jon Brenas and Lee Hart from Wellcome Sanger, for their assistance in reviewing the *Adir1* and *Amin1* API code.

## Supporting Material

**Figure S1:**
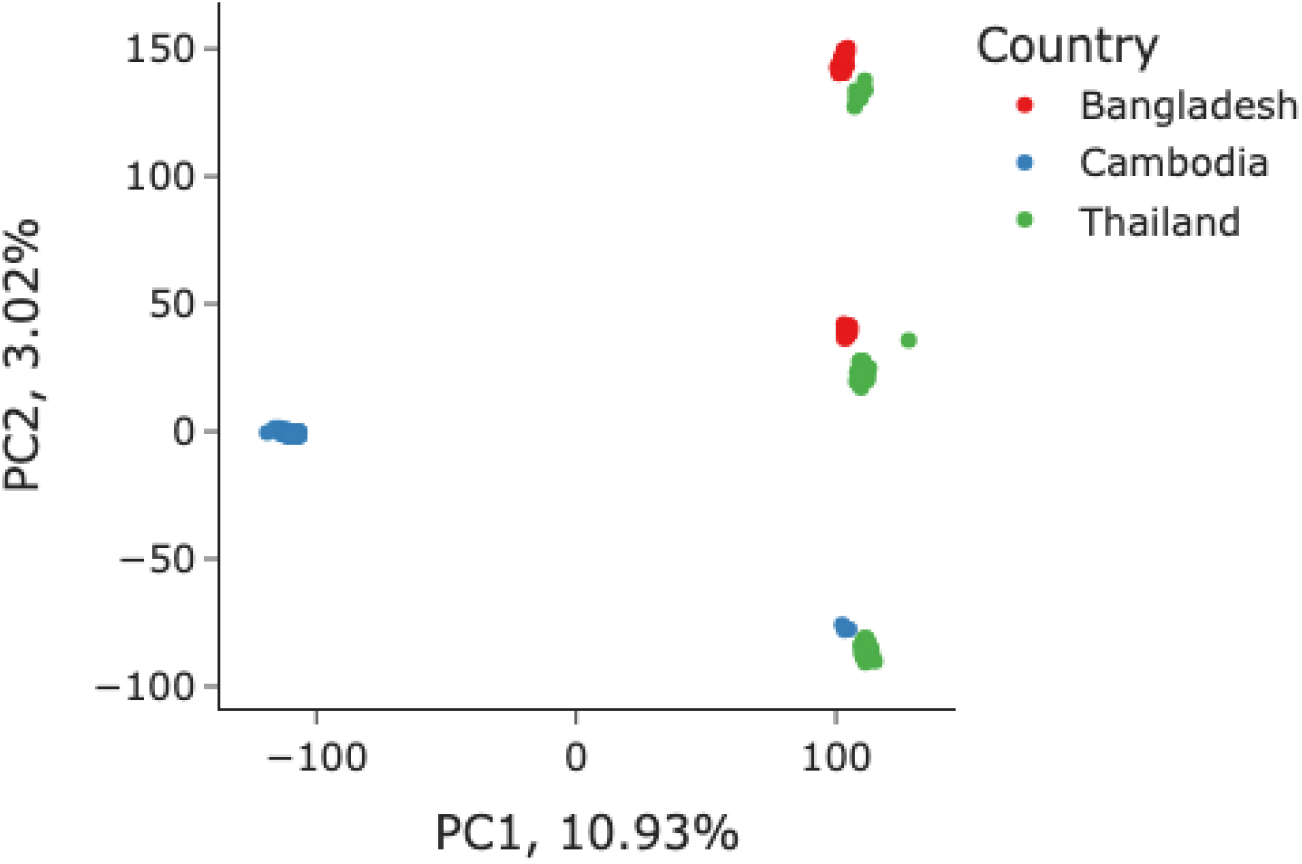
Principal component analysis (PCA) of *An. dirus* s.l. Based on 50,000 randomly chosen variants from contig KB673090. Point colour indicates country of sampling. X and Y axis indicate PC1 and PC2, with the % denoting the explained variance ratio (EVR) of the respective PC. The group on the left consists of *An dirus* samples from Cambodia, the left, *An. baimaii*. Note the clustering pattern along PC2 (y axis) suggestive of inversion polymorphism dominating the signal.

**Supplementary Tables (for in-detail column descriptions, see per-sheet legends**). **Table S1:** Per-sample metadata.

**Table S2:** Genes contained in outlier (>0.99 percentile) G123 regions in *An. dirus*.

**Table S3:** Possible resistance-associated genes contained in outlier (>0.99 percentile) G123 regions in *An. dirus*.

**Table S4:** Non-synonymous mutation frequencies in orthologs of *An. gambiae* target-site resistance genes (*Vgsc, Ace1*).

